# A comparative study of circulating tumor cell isolation and enumeration technologies in lung cancer

**DOI:** 10.1101/2024.02.05.578972

**Authors:** Volga M Saini, Ezgi Oner, Mark Ward, Sinead Hurley, Brian David Henderson, Faye Lewis, Stephen P Finn, John O’Leary, Sharon O’Toole, Lorraine O’Driscoll, Kathy Gately

**Affiliations:** Thoracic Oncology Research Group, Trinity Translational Medicine Institute, St James’s Hospital, Dublin, Ireland; Department of Clinical Medicine, School of Medicine, Trinity College Dublin, Ireland; Trinity St. James’s Cancer Institute, Trinity College Dublin, Ireland; Department of Obstetrics and Gynaecology, School of Medicine, Trinity College Dublin, Ireland; Department of Histopathology and Morbid Anatomy, School of Medicine, Trinity College Dublin, Ireland; School of Pharmacy and Pharmaceutical Sciences, Trinity College Dublin, Ireland; Trinity Biomedical Sciences Institute, Trinity College Dublin, Ireland

**Keywords:** Circulating tumor cells (CTCs), non-small cell lung cancer, CellMag™, EasySep™, RosetteSep™, Parsortix®

## Abstract

Circulating tumor cells (CTCs) have potential as diagnostic, prognostic and predictive biomarkers in solid tumors. Despite FDA approval of CTC devices in various cancers, their rarity and limited comparison between analysis methods hinder their clinical integration for lung cancer. This study aimed to evaluate five CTC isolation technologies using a standardized spike-in protocol: the CellMag™ (EpCAM-based enrichment), EasySep™ and RosetteSep™ (blood cell depletion), and the Parsortix® PR1 and next generation Parsortix® Plus (PX+) (size-based enrichment). The Parsortix® systems were also evaluated for any difference in recovery rates between cell harvest versus in-cassette staining. Healthy donor blood (5 mL) was spiked with 100 fluorescently labeled H1975 lung adenocarcinoma cell line, processed through each system and the isolation efficiency was calculated. All tested systems yielded discordant recovery rates with the CellMag™ having the highest mean recovery (70 ± 14%) followed by the PR1 (in-cassette staining) with a recovery of 49 ± 2% while the EasySep™ had the lowest recovery (18 ± 8%). The CellMag™ and Parsortix® PR1 may have potential clinical applications for lung cancer patients, albeit needing further optimization and validation.

## 1. Introduction

Circulating tumor cells (CTCs) are shed into the blood from either primary or metastatic tumors and can provide an important snapshot of tumor heterogeneity and metastatic potential. Recent advances in the sensitivity of cellular and molecular detection technologies have increased our understanding of the molecular characteristics and timing of dissemination of CTCs during cancer progression. Single-cell multi-omics technologies have shown that CTCs are heterogeneous, disseminate early and represent a subpopulation of the primary tumor responsible for disease relapse [1]. CTC enumeration has been widely used as a prognostic biomarker to monitor treatment response and predict disease relapse. In non-small cell lung cancer (NSCLC), CTC counts are low but have been shown to be prognostic in early-stage disease [2–4]. Interestingly, several studies have shown that blood collected from the pulmonary vein during surgery contains higher numbers of CTCs than blood from the peripheral vein [5–9]. Detection rates in metastatic NSCLC can be as low as 1–10 CTC/mL blood with slightly higher rates in small cell lung cancer (SCLC) [10]. In advanced disease, changes in CTC counts in response to therapy have highlighted their potential as a predictive biomarker [11, 12].

Despite the presence of CTCs indicating a poor prognosis in patients with NSCLC, their scarcity in blood compared to other cell types has hindered the translation of CTC molecular profiling to the clinic. This is a significant challenge particularly in early-stage disease where their identification and characterization could offer the most benefit for patients, helping to guide treatment and disease management. However, the field of CTC isolation is a rapidly evolving one and over the past decade there has been a surge in new CTC isolation devices and downstream workflows for molecular profiling. Current technologies are primarily based on immunoaffinity (positive or negative enrichment strategies) or biophysical properties. Although two devices are Food and Drug Administration (FDA) approved for CTC enrichment, neither are approved in NSCLC. The CellSearch® CTC Test (Menarini-Silicon Biosystems, Bologna, Italy) is approved in breast, colorectal and prostate cancer [13–15]. In 2022, the Parsortix® PC1 System (ANGLE plc, Guildford, UK) received FDA clearance for capturing and harvesting of CTCs from the blood of metastatic breast cancer patients enabling subsequent user-validated analysis [16].

Currently the assessment of CTCs has not translated into the routine clinical management of NSCLC patients. Reproducibility and sensitivity remain a challenge as do issues with manual processes and low recovery rates [17–20]. Here we evaluate the recovery rates of 5 different CTC isolation systems including: the CellMag™ system which uses a manual immuno-magnetic (ferrofluid technology) enrichment of epithelial cells using the epithelial cellular adhesion molecule (EpCAM) and is based on the same technology as the CellSearch®; EasySep™ targets cells for either removal (negative selection via CD45 depletion) or positive selection using antibody complexes directed to specific cell surface antigens; RosetteSep™ is an immunodensity procedure that isolates unlabeled CTCs using an antibody cocktail to deplete monocytes along with erythrocytes from whole blood; Parsortix® PR1 enables the separation and capture of cells present in blood based on their size and deformability [17] and the Parsortix® PX+ which is ANGLE plc’s next generation system currently undergoing beta-testing and utilizes the same isolation method as the PR1.

## 2. Materials and methods

### 2.1 Preparation of cells prior to spiking

#### 2.1.1 Cell culture

The NSCLC cell line NCI-H1975 (RRID:CVCL_1511) was used for spike-in experiments. H1975 cells are a lung adenocarcinoma cell line that express epithelial markers including EpCAM and cytokeratins [21, 22]. A variant of this cell line expressing green fluorescent protein (H1975-GFP) and stained with the CellTracker™ Green CMFDA (5-chloromethylfluorescein diacetate) dye was also utilized to ensure the detectability of all cells. This GFP transfected variant of the H1975 cell line was a kind gift from Dr. Steven Gray from Trinity Translational Medicine Institute, St James’s Hospital, Ireland. All cell lines were authenticated using short tandem repeat (STR) analysis within the past 3 years and regularly checked for the absence of mycoplasma. Both cell lines were grown in the following growth medium: RPMI 1640 + GlutaMAX™ (Gibco) supplemented with 10% fetal bovine serum (FBS) (Gibco) and 1% penicillin-streptomycin (Gibco) in a humidified atmosphere at 37°C and 5% CO2.

#### 2.1.2 Reagents and solutions

- <u>0.5 M Ethylenediaminetetraacetic acid (EDTA) (50 mL):</u> 9.305 g EDTA (disodium salt, molecular weight = 372.4 g/mol) and 40 mL deionized water. Adjust pH to 8 by adding sodium hydroxide. Make up to 50 mL with deionized water.
- <u>Spike in buffer (10 mL):</u> 40 µL 0.5 M EDTA, 2 mL 5% BSA and 8 mL DPBS without Ca^2+^ and Mg^2+^ (Cytiva).
- <u>EasySep™ Buffer (50 mL):</u> 49 mL DPBS without Ca^2+^ and Mg^2+^, 1 mL FBS and 100 µL 0.5 M (500 mM) EDTA.
- Recommended medium for RosetteSep™ (DPBS with 2% FBS, 500 mL): Add 10 mL FBS to 490 mL DPBS.

#### 2.1.3 Cell labelling using CellTracker™ dye

H1975 and H1975-GFP cells were pre-labeled with the CellTracker™ Green CMFDA dye (Catalog #C7025, Thermo Fisher Scientific, Eugene, USA) prior to spike-in experiments to facilitate detection and enumeration. Cells were at approximately 80-90% confluence on the day of staining. After trypsinization and centrifugation, the H1975 cell pellet was resuspended in 4 mL of growth media. 2 mL of this cell suspension was transferred to another 15 mL tube and centrifuged at 400 xg for 3 min. The below steps were then performed with the light off in a sterile biosafety cabinet. The cell pellet was then resuspended in 2 mL RPMI 1640 + GlutaMAX™ (Gibco) containing 2 µL of 5 mM the CellTracker™ Green CMFDA dye stock solution (1:1000 dilution, final concentration: 5 µM) and 4 µL of 1 mg/mL Hoechst 33342 dye (1:500 dilution) (Life technologies, Eugene, OR, USA) and resuspended thoroughly using a low binding pipette tip. Precautions were taken to protect the now-fluorescently labeled H1975 cells from the light. The stained cells (in 15 mL tube) were covered with tinfoil and placed in a humidified incubator at 37°C with 5% CO2 for 30 min. After incubation, the stained cell suspension was centrifuged at 400 xg for 3 min and supernatant removed. The labeled cells were washed by resuspending the cell pellet in 2 mL DPBS and then centrifuged at 400 xg for 3 min. The supernatant was removed, and the cell pellet was resuspended in 1 mL of spike-in buffer. Spike-in buffer was made fresh on the day of every experiment.

#### 2.1.4 Serial dilution of cell suspension

Cell counting was performed using trypan blue staining. Serial dilution was then performed using spike-in buffer to create a cell suspension with a final concentration of approx. 1000 cells/mL (Fig. 1). This suspension was stored in a humidified incubator at 37°C with 5% CO2 until further use (max. 15-20 min). Only low binding pipette tips and Eppendorf tubes were used in this process to minimize cell loss.

**Fig. 1.**
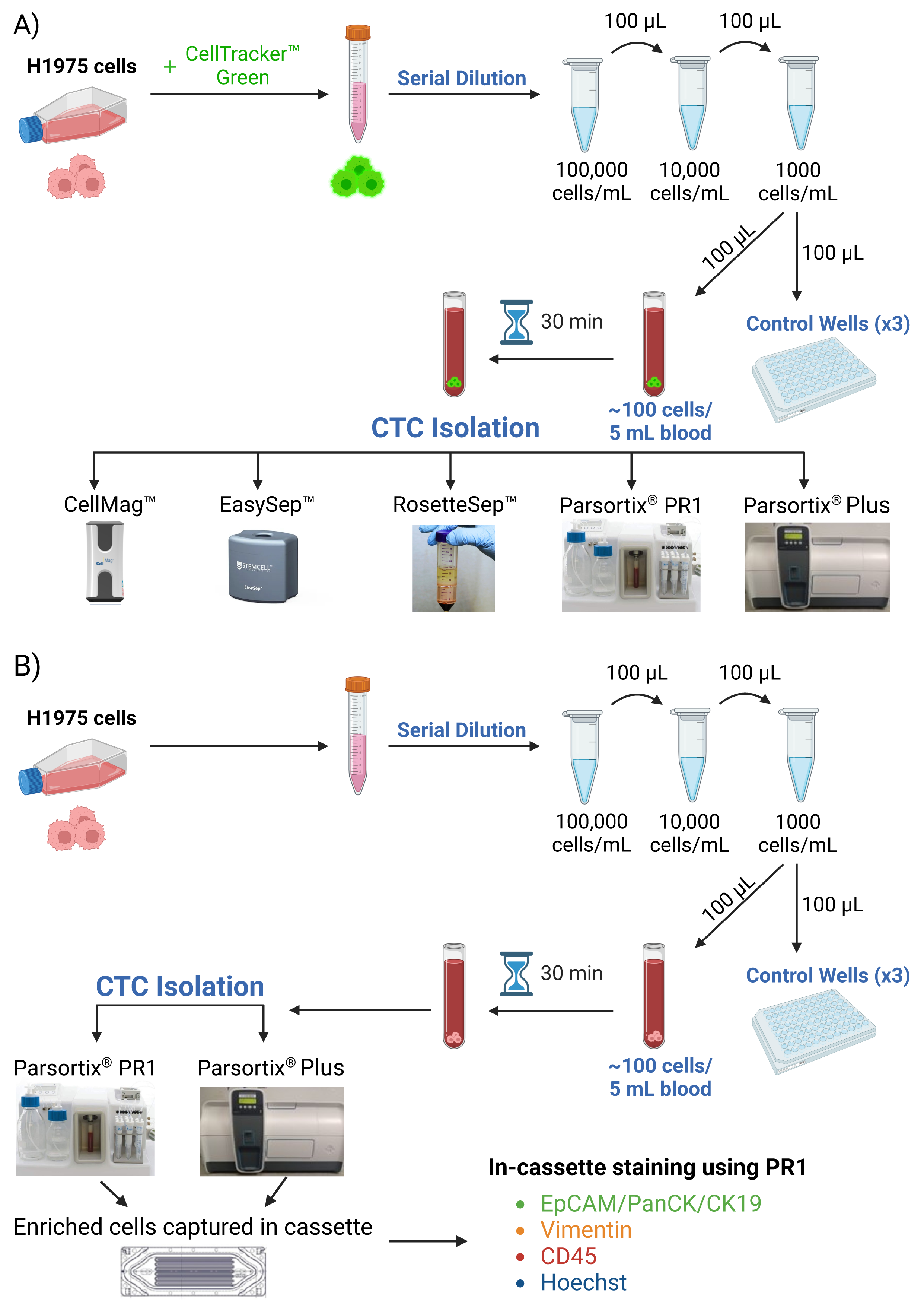
Overview of spike-in experiment and comparison of CTC isolation systems. A) H1975 or H1975-GFP cells were labeled with CellTracker™ Green CMFDA dye, and a cell suspension of 1000 cells/mL was made. 100 µL of this fluorescently labeled H1975 cell suspension (containing approx. 100 cells) were added to both 5 mL of healthy donor blood and each control well (total 3). The spiked blood sample was incubated for 30 min before processing through the different CTC isolation systems: CellMag™, EasySep™, RosetteSep™, Parsortix® PR1 and PX+. **B)** In order to compare the recovery rate of in-cassette staining between the PR1 and PX+ systems, a similar protocol was followed using unlabeled H1975 cells. The enriched cells captured in the cassette were fixed and stained using the Parsortix® PR1 with Alexa Fluor (AF)-conjugated antibodies against Cytokeratins (CK 4,5,6,8,10,13, 18 and 19) and EpCAM (AF488, green), Vimentin (AF546, yellow), CD45 (AF647, red), and Hoechst nuclear stain (blue).

#### 2.1.5 Control wells

The serial dilution process described above is prone to error and it is difficult to determine exactly how many cells are being spiked into the blood. To overcome this issue, we included 3 ‘control wells’ to estimate how many cells were spiked (on average) into blood before processing through each CTC isolation technology. In each spike-in experiment (Fig. 1), 100 µL of the cell suspension (approximately containing 100 cells) was pipetted into the blood sample and into 3 control wells. The average number of cells counted from these 3 control wells was estimated to be the approximate number of cells spiked into the blood before CTC isolation and was used to calculate the recovery rate for each system (section 2.3).

For the control wells and other steps which included cell harvest, we used wells (96-well plate) coated with poly-L-lysine (Sigma-Aldrich, Life Science, Catalog #P4707) to improve cell adhesion. Each well was coated with 50 µL of poly-L-lysine and left to incubate for 5 min with the lid on at room temperature. The poly-L-lysine was then removed, and the wells were allowed to dry (no lid) for 30 min. 100 µL of the cell suspension (approx. concentration was 1000 cells/mL) was then pipetted (using a low binding pipette tip) into 3 separate lysine coated wells (3 control wells). The 96-well plate was then placed in the incubator (37°C and 5% CO2) for 1 hour to allow the cells to settle before visualization.

### 2.2 Enrichment of spiked cells using different CTC isolation systems

#### 2.2.1 CellMag™ Epithelial CTC Kit

This semi-automated system developed by Menarini Silicon Biosystems utilizes the same method of CTC isolation as the FDA approved CellSearch® system, however, the CellMag™ system is for research use only. This technology isolates circulating epithelial cells from whole blood by immunomagnetic positive enrichment. The CellMag™ Epithelial CTC Kit consists of a ferrofluid based capture reagent that contains magnetic particles which are coated with anti-EpCAM antibodies to capture epithelial cells in circulation. After enrichment, the captured cells are labeled with fluorescent reagents for identification. These fluorescent staining reagents include DAPI (nuclear stain), anti-CK-PE (mouse monoclonal antibodies specific to cytokeratin’s conjugated to phycoerythrin) which stains epithelial cells and anti-CD45-APC (anti-mouse CD45 monoclonal antibody conjugated to allophycocyanin) which stains leukocytes (Fig. 4). In summary, the CellMag™ assay detects CTCs that are EpCAM+, CK+ and CD45-.

The CellMag™ Epithelial CTC Kit (REF: 9603) and the CellMag™ (magnet) were used to isolate spiked cells according to the manufacturers protocol (Fig. 1A). The only adjustment to the kit’s protocol was that we included an additional centrifugation step after vortexing to obtain a more enriched population. The detailed protocol is listed in Appendix 1. Healthy donor blood was collected in a CellSave Preservative tube. After precoating a serological pipette with 1% BSA, 5 mL of this blood was transferred to a conical tube (containing spiked-in H1975 cells stained with CellTracker™) and left to incubate at room temperature for 30 min before being processed through the CellMag™ system. Briefly, the sample was centrifuged at 800 xg for 10 min for plasma separation. After plasma removal, 150 µL of capture enhancement reagent and anti-EpCAM ferrofluid were added. Magnetic separation was then performed (placing the sample in CellMag™ magnet) in order to isolate the target cells. The negative fraction of the sample was aspirated using a semi-automated syringe tool. The target cells were washed again, and the supernatant was aspirated using the syringe as described above. The enriched target cells were then permeabilized and stained using the reagents included in the kit. The target cells were washed again, the fluid and unlabeled cells were removed by aspiration. Lastly, the enriched target cells were fixed, resuspended in CellMag™ buffer and transferred to one well of the 96-well plate for incubation then visualization.

#### 2.2.2 EasySep™ Direct Human CTC Enrichment Kit

The EasySep™ Direct Human CTC Enrichment Kit (Catalog #19657, StemCell Technologies) along with the “Big Easy” Magnet (Catalog #18001, StemCell Technologies) was used in this study to enrich cancer cells spiked into whole blood. This system enriches CTCs by immunomagnetic negative selection. This kit contains an antibody cocktail that targets surface markers (CD2, CD14, CD16, CD19, CD45, CD61, CD66b, and Glycophorin A) on hematopoietic cells and platelets for depletion. These unwanted cells which are labeled with antibodies and Direct RapidSpheres™, can then be separated using an EasySep™ magnet.

The spiking of cultured cells into blood was performed as above and the isolation was performed as per the kits’ instructions. The detailed protocol is listed in Appendix 2. We made various adjustments to the manufacturer’s protocol based on the protocol used by Fankhauser *et al.* to attain a highly pure enriched population [23]. Briefly, 5 mL of healthy donor blood was added to a 14 mL round bottomed tube containing the fluorescently labeled H1975 cells. Then 250 µL of enrichment cocktail (provided in kit) was added and mixed slowly by inversion 10 times, then incubated for 5 min at room temperature. RapidSpheres™ were vortexed for 30 seconds before adding 250 µL of it to the blood and then mixing it by inversion 10 times. EasySep™ Buffer (section 2.1.2) was added to double the volume and then mixed by inversion 10 times. The sample was placed in the “Big Easy” magnet and incubated at room temperature for 10 minutes. The enriched cell suspension was poured into a new 14 mL round bottomed tube and 250 µL of RapidSpheres™ was again added to the sample and mixed by inversion 10 times. The tube was again placed in the Big Easy magnet and incubated at room temperature for 10 min. The entire enriched cell suspension was pipetted into a new 15 mL conical tube. The sample was centrifuged at 800 xg for 5 min to pellet cells. The supernatant was aspirated, and the pellet was resuspended in 200 µL of DPBS containing 0.2 µL of 1 mg/mL Hoechst 33342 dye and transferred to a poly-L-lysine-coated well of a 96-well plate for incubation then visualization. Hoechst dye was added to the enriched cell suspension in order to observe the level of leukocyte contamination.

#### 2.2.3 RosetteSep™ Human CD45 Depletion Cocktail

The RosetteSep™ Human CD45 Depletion Cocktail (Catalog #15112, StemCell Technologies) was used in this study to enrich cancer cells spiked into whole blood. This kit also contains an antibody cocktail (consisting of tetrameric antibody complexes recognizing CD45, CD66b and glycophorin A) which crosslinks unwanted (hematopoietic) cells to red blood cells to form immunorosettes. These immunorosettes can be pelleted by centrifugation over a density gradient medium (has a density of 1.077 g/mL) leaving enriched cells at the interface.

The spiked in H1975 cells stained with CellTracker™ were isolated from the blood by following the kit’s instructions (Appendix 3). Briefly, 250 µL of RosetteSep™ Cocktail was added to the blood sample containing spiked cells and thoroughly mixed by pipetting and inverting the tube. The tube was then incubated at room temperature for 10 min. The blood sample was then diluted with recommended medium (section 2.1.2) and mixed gently. 15 mL of Ficoll-Paque™ Plus (#17-1440- 03, GE Healthcare,) was dispensed to a 50 mL SepMate™ (#85450, Stemcell Technologies,) tube. The diluted blood sample was slowly pipetted on top of the Ficoll in the 50 mL tube and centrifuged at 1200 xg for 10 min with the brake on. The supernatant was poured into a new standard tube and topped up with recommended medium to wash the enriched cells. The suspension was centrifuged at 300 xg for 10 min with the brake off. The supernatant was discarded by aspiration and the cell pellet was resuspended in 200 µL of DPBS and 0.2 µL of 1 mg/mL Hoechst dye (1:1000 dilution) and transferred to a poly-L-lysine-coated well of a 96-well plate for incubation then visualization.

#### 2.2.4 Parsortix® PR1

It is well established in the literature that cancer cells are more deformable and elastic, and this property could be used to isolate them from blood cells [24, 25]. The Parsortix® PR1 Cell Separation system isolates CTCs based on their size and deformability. The system consists of the instrument and a disposable separation cassette. This cassette contains steps that captures cells greater than 6.5 µm, well above the average diameter of H1975 cells, which is 17.4 µm [26]. The captured cells can then be fixed, permeabilized and stained with immunofluorescence markers (in-cassette staining) and viewed under a fluorescent microscope. Alternatively, the captured cells can be flushed out and collected (cell harvest) for downstream applications.

The spiked in cells were recovered from the blood sample as per the manufacture’s protocol (Appendix 4). Briefly, the ‘PX2_PF’ protocol was run to ‘prime’ the cassette. During this process, the cassette is filled with buffer solution, and this prepares the instrument to process a blood sample. The PX2_S99F protocol was then run for blood processing and to capture the spiked in cells in the cassette.

##### 2.2.4.1 PR1 in-cassette staining

In-cassette staining was then performed to fix and stain the captured cells in the cassette using the ‘PX2_Stain 3’ protocol (Fig. 1B). We developed an optimized antibody recipe for this staining protocol:

- <u>Fixative:</u> 360 µL DPBS and 40 µL formaldehyde solution (Sigma-Aldrich, F8775).
- <u>Antibody cocktail:</u> 3.5 µL CD45 antibody (1:120 dilution) (35-Z6, Alexa Fluor 647, SantaCruz Biotechnology, sc-1178), 3.5 µL pan-Cytokeratin antibody (1:120 dilution) (C11, Alexa Fluor 488, Biotechnology, sc-8018) (detects cytokeratins 4, 5, 6, 8, 10, 13 and 18), 3.5 µl Cytokeratin 19 antibody (1:120 dilution) (A53-B/A2, Alexa Fluor 488, SantaCruz Biotechnology, sc-6278), 3.5 µL EpCAM antibody (1:120 dilution) (HEA125, Alexa Fluor 488, SantaCruz Biotechnology, sc-59906), 3.5 µL Vimentin antibody (1:120 dilution) (V9, Alexa Fluor 546, SantaCruz Biotechnology sc-6260), 3.5 µL 1 mg/mL Hoechst 33342 dye and 399 µL of permeabilization buffer Inside Perm (#130-090-477, Inside stain Kit, MACS Miltenyi Biotec).

##### 2.2.4.2 PR1 cell harvest

The cell harvesting procedure allows pulses of liquid to be passed through the cassette in a reverse direction. This liquid detaches the captured cells and pushes them through the harvest line which can then be collected in a collection tube. Before performing the cell harvest, we performed a ‘pre-harvest flush’ (by running PX2_CT2 protocol) of the instrument which is recommended to minimize residual blood cells. After this process, the PX2_H protocol was run to harvest the captured cells. The recovered cells were harvested into a low binding Eppendorf tube in a volume of approximately 200 µL. Then, 0.2 µL of 1 mg/mL of Hoechst dye was added to this tube, and its contents were transferred to a poly-L-lysine-coated well of a 96-well plate for incubation then visualization.

#### 2.2.5 Parsortix® Plus

The Parsortix® Plus (PX+) is an upgraded version of the Parsortix® PR1 that we were beta-testing in our lab. The main differences between both systems are that PX+ is a closed system (Fig. 1) and has a faster blood processing time than the PR1 (Table 1). However, the PX+ is unable to perform in-cassette staining (only cell harvest).

**Table 1.** Summary of different circulating tumor cell isolation methods tested.

|  | **CellMag™** | **EasySep™** | **RosetteSep™** | **PR1** | **PX+** |
| --- | --- | --- | --- | --- | --- |
| **Manufacturer** | Menarini Silicon Biosystems | Stemcell Technologies | Stemcell Technologies | ANGLE plc | ANGLE plc |
| **CTC isolation/detection method** | Immunomagnetic , EpCAM dependent. | Immunomagnetic , Negative Selection via CD45 depletion. | Immunodensity , Negative Selection via CD45 depletion. | Size based microfluidic system. | Size based microfluidic system. |
| **Skill/Complexity** | Moderate | Low | Moderate | Low | Low |
| **Advantages** | Cost-effective and straightforward protocol, semi-automated. | Fast and easy to use CTC isolation process. | Results in unlabeled viable cells suitable for CTC culture. | Recovery of heterogeneous population of CTCs and ability to capture CTC clusters. | Recovery of heterogeneous population of CTCs and ability to capture CTC clusters. |
| **Disadvantages** | Only captures CTCs that express EpCAM. Time consuming and since cells are fixed, not suitable for culture. | Variable recovery rates reported and may inadvertently remove CTCs in clusters with blood cells. | May inadvertently remove CTCs in clusters with blood cells. | Only capturing CTCs above a certain size. It is time consuming to analyze stained slides. | No option for in-cassette staining. |
| **‘Hands-on-time’ for each method** | 5 hours | 2 hours | 2.5 hours | 45 min | 35 min |
| **Total Time required CTC method (isolation to enumeration)** | 6.5 hours | 3 hours | 3.5 hours | Harvest: 2 hours<br>In-cassette stain: 5 hours | Harvest: 1.25 hours<br>In-cassette stain: 3.45 hours |

##### 2.2.5.1 PX+ cell harvest

The spiked in cells were recovered from the blood sample by running the ‘PXP_SMP_v1’ protocol on the PX+ (Appendix 5.1). Like the PR1 cell harvest, the captured cells were harvested into a low binding Eppendorf tube in a volume of approximately 200 µL. Then, 0.2 µL of 1 mg/mL of Hoechst dye was added to this tube and its contents were transferred to a poly-L-lysine-coated well of a 96-well plate for incubation then visualization.

##### 2.2.5.2 PX+ in-cassette staining

The PX+ does not enable in-cassette staining. However, there is an option to remove the cassette from the PX+ before proceeding to the cell harvest. At this step (see Appendix 5.2), the casette was removed from the PX+ and in-casette staining was performed on the PR1 as described in Section 2.2.4.1 (Fig. 1B).

### 2.3 Calculation of recovery rate from CTC isolation technologies

#### 2.3.1 Enumeration of recovered cells

After the 1-hour incubation, the recovered cells (in the 96-well plate) from the CellMag™, EasySep™, RosetteSep™, PR1 and PX+ (harvest) systems and their respective control wells were imaged using an automated imaging protocol and the BioTek Lionheart FX automated microscope. We established a protocol (Gen5 3.12 software) which can take images (tiles) from all areas of the well at a magnification of 4X (Fig. 2A, Fig. S1). In our protocol, we divided the well into 6x4 image ‘tiles’ which were then ‘stitched together’ to create a ‘montage’ of the entire well. These images were taken using a Z-stack (10 focal planes) so spiked in cells at different planes of focus could still be visualized and enumerated. This protocol was set up to take images with the DAPI (Hoechst), GFP (CellTracker™ Green) and Bright Field channels. After capturing the ‘montage’ image, the number of recovered cancer cells in each individual tile was manually counted. Cells that were positive for CellTracker™ Green and Hoechst, with a morphologically intact nucleus, were counted as our recovered cancer cells (Fig. 5). For visualizing the CellMag™ harvest, the RFP and Cy5 channels were also included when ‘scanning’ the whole well to determine expression of cytokeratin and CD45 as this was part of the kit’s staining protocol (Fig. 4).

**Fig. 2.**
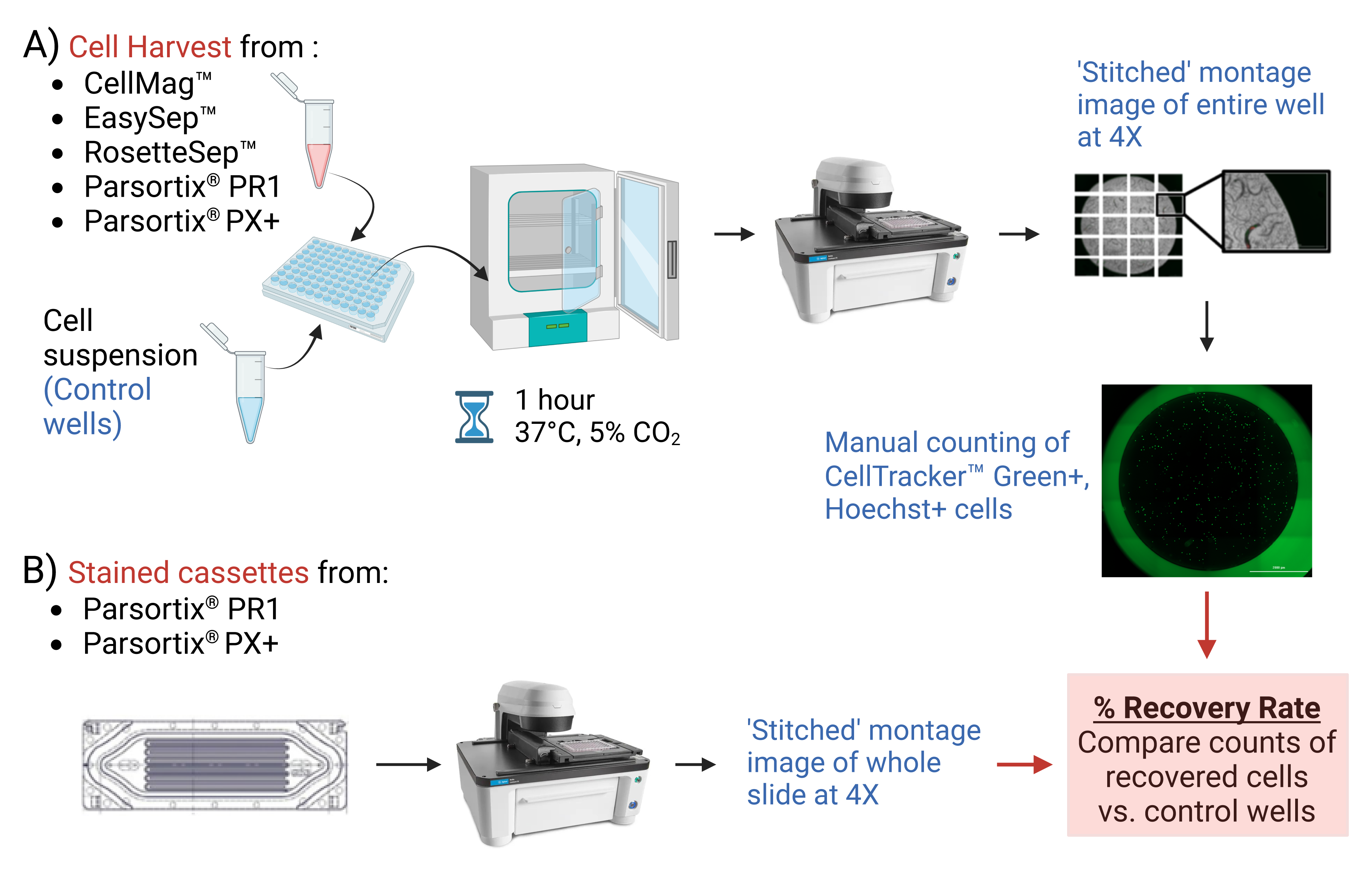
Calculation of recovery rate from CTC isolation technologies. A) After CTC isolation, the recovered cells from the CellMag™, EasySep™, RosetteSep™, PR1 and PX+ were incubated for 1 hour along with the associated control wells. Then, these wells were scanned using an imaging protocol (BioTek Lionheart FX automated microscope) to enumerate the total number of recovered cancer cells stained with Hoechst (DAPI channel) and CellTracker™ Green dye (GFP channel). **B)** The PR1 and PX+ stained cassettes were visualized using another automated imaging protocol. The percentage recovery rate was calculated by comparing the count of recovered cells against the average cell count from the control wells.

**Fig. 3.**
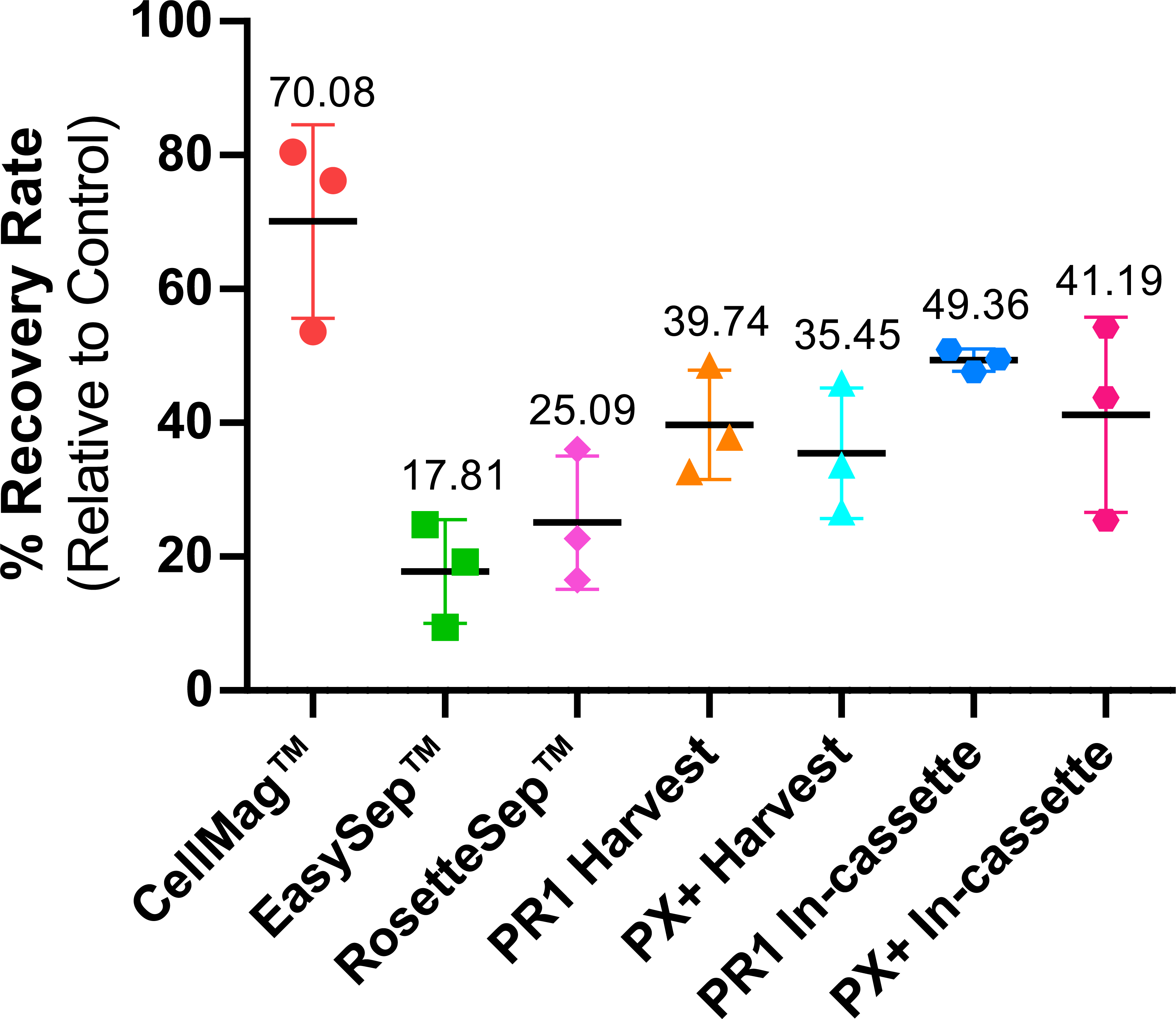
Comparison of the various methods used for CTC isolation. The lines indicate the range of the percentage recovery rates (%RR) with standard deviation (error bars) observed in three independent spike-in experiments for H1975 cells (n=3). The mean % RR for each technology is also displayed. One-way ANOVA and Tukey’s multiple comparisons test were used to compare recoveries between the 7 different isolation methods using GraphPad prism (version 8.0.2). Significant differences in %RR are represented as follows: * p<0.05, ** p < 0.01, *** p<0.001.

**Fig. 4.**
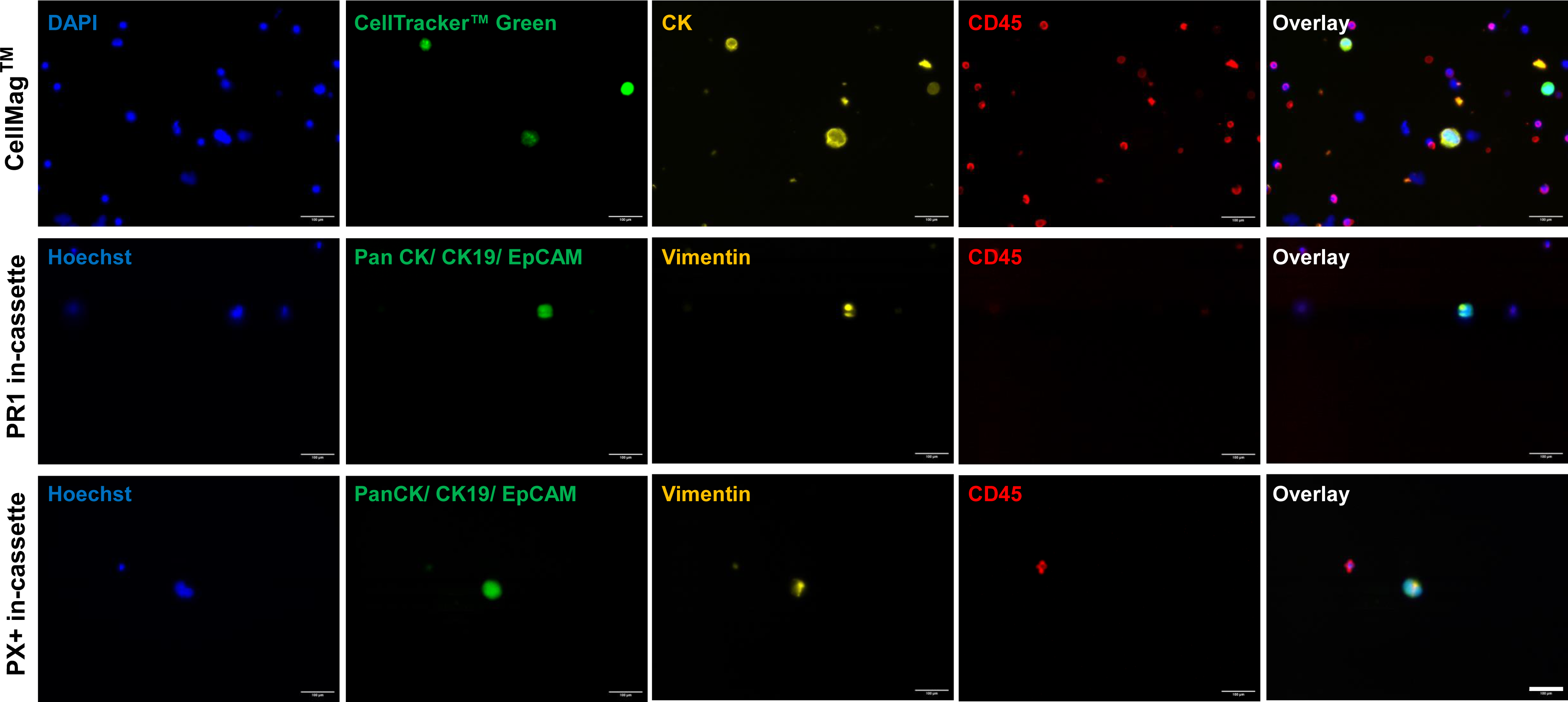
Representative staining of spiked H1975 cells enriched using different methods. Top Panel: Recovered pre-labeled (CellTracker™ Green) H1975 cells from the CellMag™ system. As per the kit’s protocol, recovered cells were stained with antibodies against cytokeratins (conjugated to phycoerythrin, yellow), CD45 (conjugated to allophycocyanin, red) and DAPI nuclear stain (blue). Middle and Bottom Panel: H1975 cells recovered using the PR1 and PX+ respectively. Captured cells were stained (in-cassette) with Alexa Fluor (AF)-conjugated antibodies against Cytokeratins and EpCAM (AF488, green), Vimentin (AF546, yellow), CD45 (AF647, red), and Hoechst nuclear stain (blue). Images were captured under 20X magnification using the BioTek Lionheart FX automated microscope and analyzed using ImageJ software. Scale bar represents 100 μm.

**Fig. 5.**
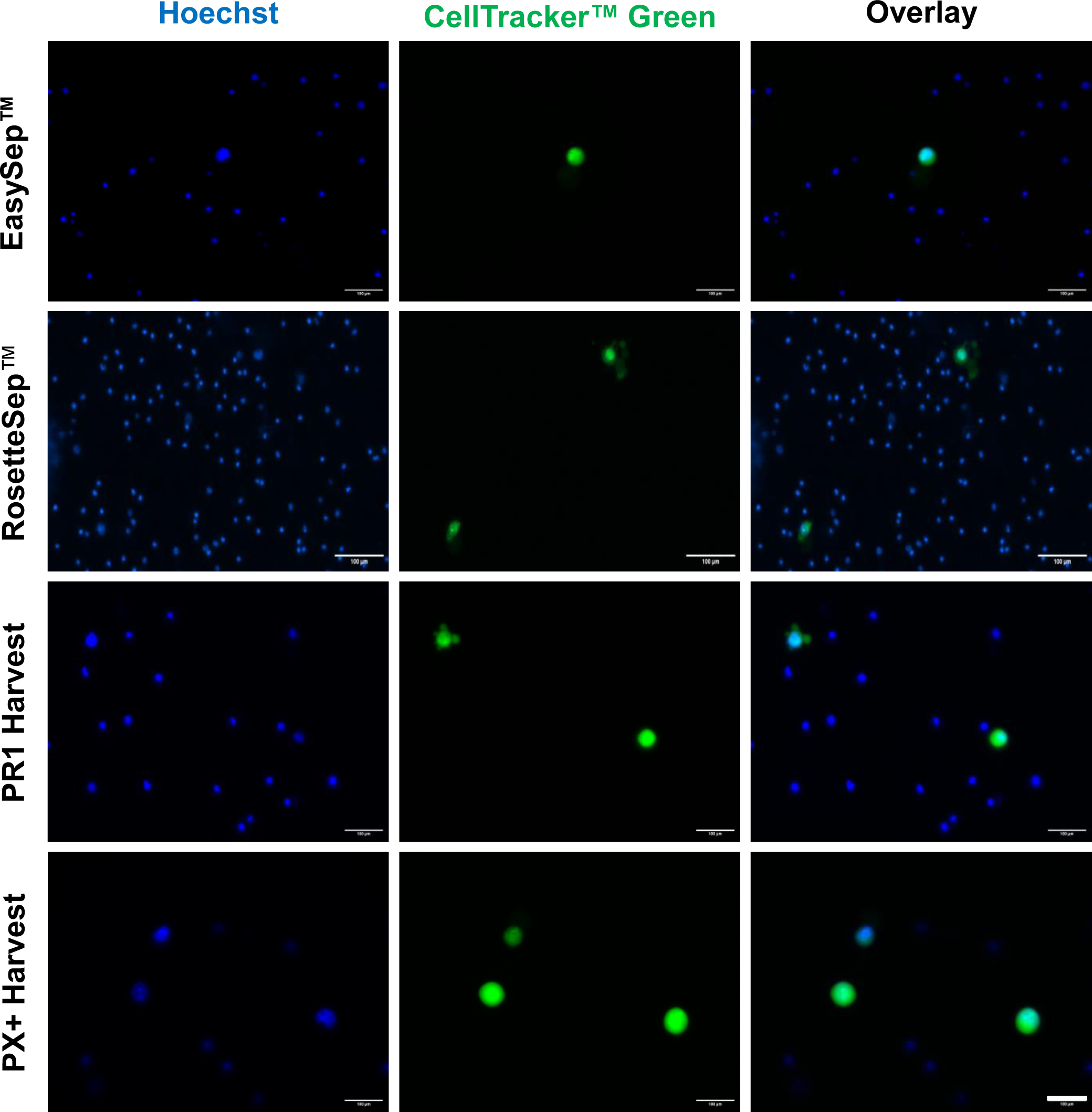
Representative images of spiked and pre-labeled H1975 cells enriched using different CTC isolation technologies. Approximately one hundred H1975 cells (pre-labeled with CellTracker™ Green) were spiked into 5 mL of whole blood and processed through the EasySep™, RosetteSep™, PR1 (harvest) and PX+ (harvest) methods. The enriched cell suspensions from each method were stained with Hoechst nuclear dye and the number of recovered cancer cells was counted to calculate recovery rate. Images were captured under 20X magnification using the BioTek Lionheart FX automated microscope and analyzed using ImageJ software. Scale bar represents 100 μm.

From the PR1 and PX+ in-cassette staining experiments, the cells captured and stained on the Parsortix® slides were visualized using a similar automated imaging protocol on the BioTek Lionheart FX automated microscope and Gen5 3.12 software (Fig. 2B, Fig. S2). We established another automated imaging protocol to scan the whole slide at 4X with the DAPI (Hoechst), GFP (CellTracker™ Green, CKs, EpCAM), RFP (Vimentin), CY5 (CD45) channels. In our protocol, we divided the slide into 12x27 (rows x columns) image ‘tiles’ which were then ‘stitched together’ to create a ‘montage’ of the entire slide. After capturing the ‘montage’ image of the whole slide, the number of recovered cancer cells in each individual tile was manually counted. Cells that were positive for Hoechst, EpCAM/PanCK/CK19 and Vimentin, and negative for CD45, with a morphologically intact nucleus, were counted as our recovered cancer cells.

#### 2.3.2 Recovery rate

The percentage recovery rate of each CTC isolation technology was calculated according to the equation below:

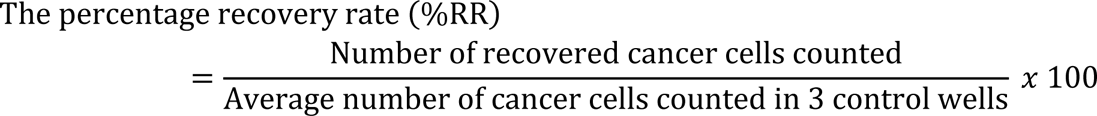

### 2.4 Statistical analysis

Three independent spike-in experiments were conducted for each method to calculate the percentage recovery rate. One-way ANOVA test and Tukey’s multiple comparisons test were used to compare the recovery rate between the 7 different isolation methods. Statistical analyses were performed using GraphPad Prism (version 8.0.2). A p value of less than 0.05 was used to indicate a significant difference.

## 3. Results

### 3.1 Recovery rates among different CTC isolation technologies

H1975, a NSCLC line was used in spike-in experiments to compare different CTC isolation technologies. The mean percentage of the recovery rates of each technology is presented in Fig. 3. The range of the percentage recovery rates (%RR) with standard deviation observed in three independent spike-in experiments for each technology is presented in Table S1.

#### 3.1.1 CellMag™

As observed in Fig. 3, the CellMag™ system had the highest recovery rate among all the methods tested with a mean of 70.08 ± 14.44 %. This method had a significantly higher rate of recovery in comparison to all the other technologies except the PR1 in-cassette staining (Table S2). This was an expected result as the CellMag™ system is based on the CellSearch® which is the gold standard for CTC isolation and FDA-approved in certain cancers. The protocol was easy to follow, and the kit comes with its own staining protocol, so no antibody optimization was needed. However, this process is very time consuming with many incubation steps, so a lot of ‘hands-on’ time is required (Table 1).

#### 3.1.2 EasySep™ and RosetteSep™

The EasySep™ and RosetteSep™ methods had the lowest recovery rates of 17.81 ± 7.73% and 25.09 ± 9.93%, respectively. Both systems from Stemcell Technologies capture CTCs by negative selection. As seen in Fig. 5, more leukocyte contamination was observed using the RosetteSep™ method in comparison to the EasySep™ method. Despite the low recovery rate, the EasySep™ system was the easiest to use and the fastest. The advantage of both methods is that they are fast and more suitable for downstream applications such as CTC culture.

#### 3.1.3 Parsortix® PR1 and PX+

The PR1 in-cassette staining yielded the second highest recovery rate (49.36 ± 1.67%). As observed in Fig. 3 (and Table S1), the spike-in results for PR1 in-cassette staining were the most consistent among the 3 replicates in comparison to all the other technologies tested. Furthermore, recovery of spiked cells from the PR1 in-cassette staining was significantly higher than the EasySep™ method (p = 0.0274). There was no significant difference in recovery rate between the PR1 and PX+ systems (in both cell harvest and in-cassette staining). This was an anticipated result as both systems utilize the same method of CTC isolation. There was a trend of increased recovery rate from in-cassette staining (both PR1 and PX+) compared to the recovery from the PR1 and PX+ harvest but these results were not significant.

## 4. Discussion

Isolating and identifying rare cells in the blood is akin to searching for a needle in a haystack. While various techniques have been developed for CTC isolation and enumeration, each technology comes with its own set of strengths and weaknesses (Table 1). Hence, comparative studies become particularly crucial, especially in scenarios where there is no FDA-approved system for CTC detection in a specific cancer type, such as the case with lung cancer.

Initial spike-in experiments with established cell lines are valuable for identifying the optimal CTC isolation and enumeration technology before handling precious patient samples. In these experiments, calculating the recovery rate based on the enumeration of cancer cells in healthy donor blood is a commonly employed analysis to assess the efficiency of these CTC technologies [27, 28, 16, 29].

The recovery rate of CTC enrichment technologies varies based on factors such as the size, morphology, and biomarker expression levels of the tested cell line, along with the compatibility of these features with the respective technique. The fact that the current gold standard, CellSearch®, is FDA-approved only for metastatic breast, colorectal, and prostate cancers underscore this variability. Since there is currently no FDA-approved CTC isolation and enumeration technology for lung cancer, we conducted a head-to-head comparison of five different technologies before working with samples from early-stage lung cancer patients.

In this study, CellMag™, the manual and ’research-use only’ version of CellSearch®, exhibited the highest recovery rate when tested with the H1975 lung adenocarcinoma cell line among other assessed technologies. To our knowledge, this is the first study comparing the isolation efficacy of CellMag™ through spike-in experiments involving cancer cells. Surprisingly, our literature review did not unveil any prior studies spiking lung cancer cells to assess the recovery rate of CellSearch®, despite its frequent utilization for CTC enumeration in lung cancer patients [30–34]. Despite observing a commendable 70% recovery of EpCAM(+) H1975 lung cancer cell line with CellMag™ in our study, its advanced version, CellSearch®, was only able to detect ≥2 CTCs in 20% of the 168 lung cancer patients, as reported by Allard *et al.* [35]. Additionally, the capture rate with CellSearch® (32%) was lower compared to Parsortix® PR1 (harvest) (61%) when analyzing blood samples from 97 NSCLC patients in parallel [36]. The primary factor contributing to this diminished sensitivity in lung cancer patient samples may be the missing of EpCAM(-) CTCs with mesenchymal characteristics, which is related to the tumor’s metastatic potential [37], shorter progression-free survival [38], and poorer survival [39] in lung cancer [40, 41]. Consequently, the exploration of marker-independent CTC enrichment technologies becomes crucial, particularly for lung cancer cases [42]. Despite exhibiting lower recovery rates in spike-in experiments, size-based Parsortix® systems or negative depletion-based RosetteSep™ and EasySep™ emerge as potential alternatives. Furthermore, the flexibility of using customizable antibodies for CTC enumeration allows for the selection of the markers of interest for each patient after enrichment with these technologies. This adaptability enhances the potential for a more personalized and effective approach in the context of lung cancer diagnostics and monitoring.

Among these technologies, Parsortix® PR1 (in-cassette staining) demonstrated the second-highest recovery rate, coupled with the highest consistency between biological replicates. While the hands-on time is minimal with semi-automated Parsortix® systems, the process of scanning the entire cassette for CTC enumeration could become time-consuming when employing a manual microscope, as discussed later. The harvest option primarily addresses imaging issues encountered with in-cassette staining and is advantageous for subsequent analyses like whole-genome sequencing (WGS) [43], single cell RNA sequencing (scRNAseq) [44, 45], PCR [44, 46, 26], fluorescence in situ hybridization (FISH) [44] and culturing CTCs [47, 48]. Moreover, it is well-suited for analyzing both circulating tumor DNA (ctDNA) and CTCs from the same blood tube [49]. However, harvest had slightly lower recovery rates compared to in-cassette staining as observed with both Parsortix® systems (p>0.05). Parsortix® PX+, the upgraded version of PR1 still under development, quickened the blood processing time and eliminated the need for flicking the blood tube to avoid unwanted clogging during enrichment. However, the recovery rate of PX+ was found slightly lower than PR1 with both harvest and in-cassette staining protocols (p>0.05) and requires PR1 for in cassette staining.

On the other hand, RosetteSep™ and EasySep™ serve as cost-effective CTC enrichment technologies by eliminating labeled blood cells using density-gradient centrifugation and magnetic fields, respectively. In this study, their recovery rates were found lower than those of CellMag™ and Parsortix® PR1 and PX+ systems when 100 cells were spiked. One possible explanation for their lower isolation efficiency could be the loss of some CTCs due to encapsulation within the unwanted blood cell clusters to be eliminated. However, instead of enumeration, these systems are particularly favorable for culturing CTCs. Especially RosetteSep™, both the human CD45 depletion cocktail and the CTC enrichment cocktail containing anti-CD36, is widely used for culturing CTCs in lung cancer [50–52] as well as other cancer types [53–55]. However, EasySep™ is generally combined with another enrichment technology for culturing CTCs. Liao *et al.* developed a two-step enrichment protocol using EasySep™^®^ human CD45 depletion kit followed by spheroid culture to isolate both epithelial and/or mesenchymal CTCs and cultured these with a success rate of 46% in blood samples of 13 head and neck cancer patients [56]. In a later study, they also added the erythrocyte lysis step before EasySep™ and spheroid culture to isolate a better-purified CTC population [57]. Interestingly, Liu *et al.* discovered the highest recovery rate using only the EasySep™ human CD45 depletion kit (58%) after spiking 100 SW620 colon cancer cells into 5 mL of blood, compared to the recovery rates of using only the EasySep™^®^ EpCAM(+) FITC-positive selection kit (25%), or a combination of both (22.5%), which were similar to our 18% recovery rate with the H1975 cell line [58]. Additionally, they successfully cultured and passaged CTCs from the pleural effusion of NSCLC patients as well as the ascetic fluids of colon and ovarian cancer patients using the depletion kit alone.

As depicted above, there is considerable variability in recovery rates observed across different studies. Interestingly, this variability persists even when the same technology and cell lines are employed. Ntzifa *et al.* spiked 10, 100, and 1000 H1975 cells into 10 mL of blood and assessed the recovery rate of Parsortix® PR1 based on Ct (cycle threshold or quantification cycle, Cq) values for CK19 mRNA expression using qPCR, rather than enumeration via immunofluorescence imaging [59]. Meanwhile, Obermayr *et al*. enhanced the calculation of recovery rate by employing both qPCR and immunofluorescence imaging techniques [26]. They introduced 100 H1975 cells into 18 mL of blood, resulting in a Ct value for CK19 of approximately ∼19, in contrast to Ntzifa *et al.*’s Ct value of 34. Moreover, they achieved an average recovery rate of 80% based on immunofluorescence, surpassing our recovery rates with both in-cassette (49%) and harvest (40%) protocols. These disparities in recovery rates among studies may stem from their enumeration protocol, which involves counting pre-stained fluorescent cells in-cassette without fixation and antibody-staining steps before harvesting them for qPCR analysis. In a comprehensive investigation, Papadaki *et al.* also spiked H1975 cells together with A549 and SK-MES lung cancer cell lines (100 cells/5 mL) to compare semi-automated approaches ISET and Parsortix® with manual enrichment methods, namely density gradient medium and erythrocyte lysis buffer, along with their combined usage with CD45 depletion technology employing beads, Dynabeads [60]. Notably, direct air drying on lysine-coated slides resulted in a higher recovery rate than cytospin (500 xg for 2 min vs. 5 min) following cell harvesting. Interestingly, their recovery rate of H1975 cells post-harvest with air-drying on lysine-coated slides (57 ± 11%) was found to be higher than our recovery rate with PR1 harvest (40 ± 8%).

There are various parameters, including spiked cell number, blood volume, type of blood collection tube, time to process blood, imaging system, researcher’s expertise especially in manual technologies, and the approach used for recovery rate calculation, that may contribute to these inconsistencies in the literature regarding the recovery rate of CTC isolation and enumeration techniques.

As there is no universally established protocol for spike-in experiments, methodologies can vary among research groups and may not always be explicitly detailed in the literature. To achieve a cell suspension with a specific cell count, researchers commonly employ the direct serial dilution method [47] or combine this with a second counting step to determine the absolute spiked cell count in the blood tube [16] or readjust cell count with further dilution by including control wells [60, 16] before introducing healthy donor blood. In our study, we chose the serial dilution method and included control wells with an equivalent volume of the spiked-in cell suspension for comparative analysis. The observed cell count in control wells varied around 100, sometimes falling below or exceeding that number. Consequently, we calculated the recovery rate as a percentage, based on the spiked cell count after isolation relative to the average cell count from these control wells. Although no significant differences were found in the %RR values, whether normalized relative to the control well or not, narrower standard deviation values were observed with %RR relative to control (Table S1). Additionally, the difference between the %RR of EasySep™ and PR1 (in-cassette staining) changed from not statistically significant (p=0.2392, Table S3) to significant (p=0.0274) when normalized (Table S2). We conclude that incorporating control wells and subsequently normalizing the recovery rate relative to these wells in the serial dilution method can enhance accuracy, particularly when spiked cell numbers are low, such as 100 cells or below.

Spiked cell number is another critical parameter that can influence the recovery rate. Drucker *et al.* conducted experiments where they spiked varying cell numbers (10, 100, 1000 and/or 10,000 cells) of the MDA-MB-231 breast cancer cell line to assess the isolation efficacy of DynaBeads^®^, EasySep™, RosetteSep™ (containing anti-CD36 system), and ScreenCell^®^ [27]. While there were no significant changes in %RR for RosetteSep™ (34-40%) when 1000, 100, and 10 cells were spiked, inter-assay variability was significant when only 10 cells were spiked. Additionally, a significant difference in %RR was observed when reducing the spiked cell number from 10,000 to 1000 with both EasySep™ and Dynabeads CD45-depletion technologies followed by an (EpCAM+) based second enrichment.

Although we fixed the spiked cell number at 100 cells for comparing five different technologies in this study, testing a wider range of spiked cell numbers may provide better insights into the detection limit value for each technique’s recovery rate.

On the other hand, some technologies have demonstrated consistent recovery rates even across a wide range of spiked cell numbers. Generally, automated technologies show higher consistency compared to manual ones. For example, harvesting with the Parsortix® PC1 system showed linearity when blood samples were spiked with 2-100 cells of live and/or fixed SKBR3, MCF7, and Hs578T breast cancer cell lines (R² ≥ 0.94) [16]. Therefore, the proximity of the spiked cell count to the CTC counts in the targeted patient population is crucial, especially when dealing with early-stage cancer patients with potentially low CTC counts, which could range from 0 to 5 [11].

Additionally, extra caution is necessary when utilizing fluorescent cell lines in spike-in experiments. It is imperative to verify the homogeneity of fluorescent intensity within the cell population by employing a fluorescent microscope or a flow cytometer before initiating any spike-in experiment, especially when utilizing ’stable’ cell lines expressing fluorescent proteins. This precaution is essential because some cells within this population, initially expressing fluorescent proteins, may lose the gene coding for the fluorescent protein while retaining the selection gene, such as an antibiotic-resistance gene, over time (i.e., in later cell passages) [61]. On the contrary, pre-staining a non-fluorescent cell line with a fluorescent dye for live cells is a commonly employed approach in spike-in experiments [26, 60]. However, it is crucial to note that the fluorescent signal can be substantially low when cells are stained in the presence of serum. Additionally, optimizing staining conditions, such as determining the optimal dye concentration and labeling time, is a prerequisite before conducting spike-in experiments. In this study, a combined approach is employed, involving the application of a fluorescent dye on the cell line expressing the fluorescent protein. This method ensures the uniformity of the cell population with respect to the fluorescent signal.

Another crucial factor influencing the CTC enumeration process is the imaging system. Whether the system is manual or automated significantly impacts the reliability of recovery rate results and should not be underestimated. Firstly, manual imaging requires heightened attention from researchers to avoid the inadvertent recounting of the same cell. On the other hand, automated imaging systems prove to be a substantial time-saver compared to their manual counterparts. For instance, in Parsortix® PR1 and PX+ systems, employing in-cassette staining can prolong the comprehensive scanning of the entire cassette using a manual fluorescent microscope. Moreover, addressing challenges arises when dealing with multiple planes within microfluidic cassettes for size-based cell selection. Similar challenges may be encountered in other sized-based CTC isolation and enumeration technologies employing filtration such as ISET, ScreenCell-Cyto, and CD-PRIME/CD-FAST Auto [62]. Consequently, these systems may require advanced imaging systems explicitly designed to automatically scan large areas quickly, operate at higher magnifications (e.g., 100X, 400X), and include Z-stack imaging options. These advanced imaging systems are generally integrated with a CTC enumeration software, which is essential to facilitate cell imaging easily and standardize the enumeration process. However, it is worth noting that open-source image analysis software, such as ACCEPT [63], specifically developed for identifying and counting CTCs, can also significantly enhance the analytical capabilities of researchers. In addition to adopting advanced imaging systems and software solutions, the recently introduced Portrait+ CTC staining kit and its customizable version, Portrait Flex, offer the flexibility to image a smaller area (comparable in size to one well of a 96-well plate) on the unique cytospin slides included in the kits post-harvesting by Parsortix® systems [64, 65].

Apart from CTC isolation technology, the recovery rate is influenced by the researchers’ familiarity with the technology, particularly in cases where manual handling is predominantly utilized. In this study, we noted enhanced recovery rates in the last replicates when employing CellMag™, EasySep™, and RosetteSep™ (Table S1). While there was no statistically significant difference in recovery rates between the semi-automated Parsortix® PR1 and PX+ systems with both harvest and in-cassette protocols (p>0.05), greater consistency in recovery rate values was observed with the commercialized Parsortix® PR1 using the in-cassette protocol.

## 5. Conclusions

In the current study, a series of manual and automated CTC isolation technologies were compared for their efficiency to enrich CTCs. Spike-in experiments using the H1975 cell line were used to evaluate the recovery rate of the following CTC isolation methods/systems: CellMag™, EasySep™, RosetteSep™, Parsortix® PR1 and PX+ systems. Each method had its own advantages and disadvantages. Direct comparison of recovery rates and provided discordant recovery rates with the CellMag™ system and Parsortix® PR1 in-cassette staining method yielding the highest recovery of cells. The EpCAM-dependent CellMag™ system had the highest mean recovery rate but the size-based Parsortix® PR1 system had the most consistent recovery rates among all the technologies tested. While further optimisation and validation are required, here we present a number of emerging technologies for the analysis of these important cells. Undoubtedly, continued industry/academy/clinical collaborators will ensure timely advancement of such technologies for the benefit of cancer patients.

## Acknowledgements

This research was supported by the North-South Research Programme administered by the Higher Education Authority on behalf of the Department of Further and Higher Education, Research, Innovation and Science and the Shared Island Fund (CLuB: The All-Ireland Cancer Liquid Biopsies Consortium (https://www.clubcancer.ie). The authors would like to thank ANGLE plc, UK for kindly providing the Parsortix® PR1 and PX+ instruments for this study. We would also like to thank Stemcell Technologies, UK for kindly providing the EasySep™ and RosetteSep™ kits used in this study. We acknowledge Dr. Steven Gray for providing the valuable H1975-GFP cell line used in the current study and all the healthy volunteers for providing the blood samples.

## Author contributions

Conceptualization, V.M.S., E.O., K.G.; Experiments, V.M.S., E.O.; Methodology, V.M.S., E.O., M.W., K.G.; Blood sample collection from healthy donors, S.H.; writing—original draft preparation, V.M.S., E.O., K.G.; writing—review and editing, V.M.S., E.O., M.W., S.F., B.D.H., F.L., J.O.L., S.O.T., L.O.D., and K.G.; funding acquisition, L.O.D., K.G., J.O.L. and S.O.T. All authors have read and agreed to the published version of the manuscript.

## Data accessibility

The data that support the findings of this study are openly available in Figshare at http://doi.org/10.6084/m9.figshare.25146299 and http://doi.org/10.6084/m9.figshare.25146248.

## Supporting Information

The data that supports the findings of this study are available online in the supplementary material of this article:

**Appendix 1.** Protocol for CellMag™ Epithelial CTC Kit

**Appendix 2**. Protocol for EasySep™ Direct Human CTC Enrichment Kit

**Appendix 3**. Protocol for RosetteSep™ Human CD45 Depletion Cocktail

**Appendix 4**. Protocol for Parsortix® PR1

**Appendix 4.1.** Protocol for Parsortix® PR1 in-cassette staining

**Appendix 4.2.** Protocol for Parsortix® PR1 cell harvest

**Appendix 5**. Protocol for Parsortix® Plus (PX+)

**Appendix 5.1.** Protocol for Parsortix® Plus (PX+) cell harvest

**Appendix 5.2.** Protocol for Parsortix® Plus (PX+) in-cassette staining

Fig. S1. Automated imaging protocol for scanning a well of 96-well plate at 4X to image cell harvest from CellMag™, EasySep™, RosetteSep™, Parsortix® PR1 and PX+ systems and the control wells.

Fig. S2. Automated imaging protocol for scanning the whole Parsortix® slides at 4X to image after Parsortix® PR1 and PX+ in-cassette staining process.

Table S1. Overview of the percentage of recovery rates from CTC isolation technologies tested in this study.

Table S2. Results of Tukey’s multiple comparisons test displaying the difference and significance in recovery rates (normalized to control).

Table S3. Results of Tukey’s multiple comparisons test displaying the difference and significance in recovery rates (not normalized to control).

