## Supplemental Appendix 1-5 for "A comparative study of circulating tumor cell isolation and enumeration technologies in lung cancer"

**Appendix 1.** Protocol for CellMag™ Epithelial CTC Kit

**Appendix 2.** Protocol for EasySep™ Direct Human CTC Enrichment Kit

**Appendix 3.** Protocol for RosetteSep™ Human CD45 Depletion Cocktail

**Appendix 4.** Protocol for Parsortix® PR1

**Appendix 4.1.** Protocol for Parsortix® PR1 in-cassette staining

**Appendix 4.2.** Protocol for Parsortix® PR1 cell harvest

**Appendix 5.** Protocol for Parsortix® Plus (PX+)

**Appendix 5.1.** Protocol for Parsortix® Plus (PX+) cell harvest

**Appendix 5.2.** Protocol for Parsortix® Plus (PX+) in-cassette staining

### Appendix 1. Protocol for CellMag™ Epithelial CTC Kit

#### 1. Sample Preparation

1) Collect about ~6 mL healthy donor blood in a CellSave preservative tube (Menarini silicon biosystems, 9598-20). After precoating a serological pipette with 1% BSA, transfer 5 mL of blood to conical tube provided and leave to incubate at room temperature in the dark for 30 min, after which it was inverted 5 times.
2) Using a new pipette add 6.5 mL of Dilution Buffer into the conical tube (containing blood with spiked in cells).
3) Close the conical tube using a conical tube cap provided and invert it five times.
4) Use a swing bucket centrifuge to centrifuge the conical tube at 800 xg for a full 10 min in the brake off mode. Analyse the tube to ensure that the red blood cell layer is separated from the plasma. Discard if there is no separation.
5) Open the conical tube and use a 10 mL pipette to aspirate the plasma. Leave about 3–4 mL of plasma and set the cap aside for the following steps.
6) Using a P1000 pipette carefully aspirate the remaining plasma. Leave at least 0.5–1 mL of plasma.
7) Add 3 mL of CellMag™ Buffer and 3 mL of Dilution Buffer.
8) Close the conical tube and invert it three times.
9) Open the conical tube and set aside the cap for the following steps.
10) Add 150 µL of Capture Enhancement Reagent.
11) Add 150 µL of Anti-EpCAM ferrofluid.
12) Close the conical tube and invert it three times.

#### 2. Magnetic Separation

1) Insert the conical tube in CellMag™ and leave it in for 10 min for magnetic separation.
2) Remove the conical tube from CellMag™ and invert it five times.
3) Repeat steps 1 and 2.
4) Remove the conical tube cap and discard it.
5) Mount the cannula guide on the conical tube.
6) Put the conical tube in CellMag™ and leave it in for 20 min. Meanwhile, prepare the provided syringe for a complete aspiration by locking the spring lock in first position.
7) At the end of the incubation, do not remove the conical tube from CellMag™.
8) Without removing the conical tube from CellMag™, insert the cannula with extension tubing in the cannula guide until it touches the bottom of the tube.
9) Rotate the spring stopper counter clockwise to release the syringe plunger and aspirate the negative fraction.
10) Remove the syringe from the syringe holder.
11) Push the spring stopper to discard the aspirated negative fraction.
12) Remove the conical tube from CellMag™ and put it in a rack.
13) Remove the spring stopper, the spring and the syringe holder and discard the syringe with cannula and extension tubing.

**3. Sample Washing**

1) Close the conical tube with a new conical tube cap and use a swing bucket centrifuge to centrifuge the conical tube at 300 xg for 1 min.
2) Open the conical tube and set aside the conical tube cap for the following steps.
3) Add 1.5 mL of CellMag™ Buffer against the conical tube wall, ensuring to completely wash the sides of the tube.
4) Repeat step 3 with 1.5 mL of Dilution Buffer. Close the conical tube.
5) Mix the conical tube with the vortex at 1600 RPM for 10 s. Centrifuge the conical tube at 1600 RPM for 10 s.
6) Open the conical tube and set aside the conical tube cap on a clean lint-free wipe for the following steps.
7) Mount a new cannula guide on the conical tube.
8) Put the conical tube in CellMag™ and leave it in for 10 min. Prepare the syringe as described in complete aspiration.
9) At the end of the incubation, do not remove the conical tube from CellMag™.
10) Insert a new cannula with extension tubing in the cannula guide until it touches the bottom of the tube.
11) Rotate the spring stopper counterclockwise to release the syringe plunger and aspirate the supernatant.
12) Push the spring stopper to discard the aspirated supernatant.
13) Set aside the cannula and the extension tubing on a clean lint-free wipe for the following steps.
14) Remove the conical tube from CellMag™.

**4. Sample permeabilization and staining**

1) Add the reagents against the conical tube wall, ensuring to completely wash the sides of the tube, and in the following order: 200 µL of Dilution Buffer, 150 µL of Permeabilization Reagent, 150 µL of Staining Reagent, 150 µL of Nucleic Acid Dye, 200 µL of CellMag™ Buffer.
2) Resuspend the sample five times by gently pipetting.
3) Close the conical tube and incubate it at room temperature for 20 min in the dark.
4) Open the conical tube and set aside the conical tube cap for the following steps.
5) Resuspend the sample five times by gently pipetting.
6) Without touching the liquid located at the bottom of the conical tube, add the reagents against the wall of the conical tube in the following order: 1 mL of CellMag™ Buffer and then 1 mL of Dilution Buffer.
7) Resuspend the sample five times by gently pipetting.
8) Mount the cannula guide on the conical tube.
9) Put the conical tube in CellMag™ and leave it in for 15 min. At the end of the incubation, do not remove the conical tube from CellMag™. Meanwhile, prepare the syringe for limited aspiration by locking the spring stopper in second position. Attach the cannula with extension tubing to the syringe and rotate the luer lock clockwise to lock it.

10) Insert the cannula with extension tubing in the cannula guide until it touches the bottom of the tube.
11) Rotate the spring stopper counterclockwise back to first position. This starts a limited aspiration that lasts approximately 20 seconds and only aspirates a small part of the sample.
12) Remove the syringe from the syringe holder.
13) Carefully extract the cannula with the extension tubing from the conical tube.
14) Remove the cannula guide, the spring stopper, the spring and the syringe holder and discard the syringe, the cannula with the extension tubing and the cannula guide.
15) Close the conical tube and use a swing bucket centrifuge to centrifuge it at 300 xg for 5 min.
16) Open the conical tube and set aside the conical tube cap for the following steps.
17) Using a P1000 pipette, aspirate the supernatant without touching the pellet. Leave around 100 µL of sample or keep the liquid meniscus at the level of the mark on the bottom of the conical tube.
18) Add 1 mL of CellMag™ buffer and resuspend the sample five times.
19) Close the conical tube and use a swing bucket centrifuge to centrifuge it at 300 xg for 5 min.
20) Open the conical tube and set aside the conical tube cap for the following steps.
21) Using a P1000 pipette, aspirate 1 mL without touching the pellet.

**5. Fixative**

1) Add the reagents in the following order to the bottom of the conical tube without touching the tube walls: 150 µL of Cell Fixative and then 100 µL of CellMag™ Buffer.
2) Close the conical tube and mix it with the vortex at 1600 RPM for 10 s. Centrifuge the conical tube at 1600 RPM for 10 s.
3) Incubate the conical tube at room temperature for 20 min in the dark.
4) Transfer this enriched cellular suspension (approx. 300 µL) to a poly-L-lysine-coated well of a 96 well plate. Incubate the plate in the dark (at 37°C and 5% $CO_2$) for 1 hr before visualization.

### Appendix 2. Protocol for EasySep™ Direct Human CTC Enrichment Kit

1) Using low-binding tip, add 100 µL of H1975 cell suspension stained with CellTracker™ Green (1000 cells/mL) into a 14 mL polystyrene round-bottom tube.
2) Collect approx. 6 mL healthy donor blood in green capped heparin tube (BD-Plymouth. PL6 7BP.UK, 8083925). The kit recommended use of heparin /Acid citrate dextrose (ACD) anticoagulant (instead of K2EDTA or K3EDTA) for optimum red blood cell depletion so green capped heparin tubes were used for blood collection.
3) After precoating a serological pipette with 1% BSA, transfer 5 mL whole blood from heparin tube into 14 mL polystyrene round-bottom tube containing spiked in cells (100 µL).

4) Let it incubate for 30 min in dark and then invert 5 times before processing.
5) Add 250 µL of Enrichment Cocktail to sample (50 µL/mL of sample).
6) Mix thoroughly by pipetting up and down with p1000 and low binding pipette tip. Invert the tube 10 times slowly and incubate at room temp for 5 minutes.
7) Vortex RapidSpheres™ for 30 seconds. Make sure it forms a uniform suspension and particles should appear evenly dispersed.
8) Add 250 µL of RapidSpheres™ (50 µL/mL of sample) to the sample and mix thoroughly by pipetting up and down with p1000 and low binding pipette tip. Invert the tube 10 times slowly.
9) Add 4.5 mL of EasySep Buffer (section 2.1.2) to top up the sample. Mix by gently pipetting up and down 2 -3 times. Invert the tube 10 times slowly.
10) Place the tube (without lid) into the magnet and incubate at room temperature for 10 minutes.
11) Pick up the magnet, and in one continuous motion invert the magnet and tube, pouring the enriched cell suspension into a new 14 mL round bottomed tube. Leave the magnet and tube inverted for 2-3 seconds then return upright.
12) Add 250 µL RapidSpheres™ to the new tube containing the enriched cells and mix by pipetting up and down with p1000 and low binding pipette tip. Invert the tube 10 times slowly.
13) Place the tube (without lid) into the magnet and incubate at room temperature for 10 minutes.
14) Pick up the magnet, and in one continuous motion invert the magnet and tube, pouring the enriched cell suspension into a new 14 mL round-bottom tube. Leave the magnet and tube inverted for 2-3 seconds then return upright.
15) Repeat steps 13 and 14 again. At the end of step 14, collect into a new 15 mL centrifuge tube (instead of 14 mL round bottomed tube).
16) The enriched cells are now ready to use (approx. 7.5 mL enriched cell suspension).
17) Centrifuge cellular suspension at 800 xg for 5 min.
18) Discard all supernatant very carefully without touching the cell pellet at the bottom of the tube.
19) Resuspend cell pellet in 200 µL of DPBS containing 0.2 µL of 1 mg/mL Hoechst 33342 dye (1:1000 dilution) using a low binding pipette tip.
20) Transfer this enriched cellular suspension to a poly-L-lysine-coated well of a 96 well plate. Incubate the plate in the dark (at 37°C and 5% $CO_2$) for 1 hr before visualization.

#### Appendix 3. Protocol for RosetteSep™ Human CD45 Depletion Cocktail

1) Collect approx. 6 mL healthy donor blood in 10 mL vacutainer coated with EDTA (BD-Plymouth. PL6 7BP.UK, 8083925).
2) Using a low retention pipette tip, add 100 µL of H1975 cell suspension stained with CellTracker™ Green (1000 cells/mL) into a new 50 mL tube.

3) After precoating serological pipette with 1% BSA, transfer 5 mL whole blood from EDTA tube into 50 mL tube containing spiked in cells.
4) Let it incubate for 30 min in dark and then invert 5 times before processing.
5) Add 250 µL of RosetteSep™ Cocktail to sample (50 µL/mL of sample).
6) Mix thoroughly by pipetting up and down with p1000 and invert tube and incubate at room temperature for 10 min.
7) Dilute sample with 5 mL RosetteSep™ recommended medium (section 2.1.2) and mix gently.
8) Dispense 15 mL of Ficoll-Paque™ Plus (GE Healthcare, 17-1440-03) very slowly to the 50 mL SepMate™ (Stemcell Technologies, 85450) tube.
9) Slowly pipette diluted blood sample on top of the Ficoll in the 50 mL tube, being careful to minimise their mixing).
10) Centrifuge at 1200 xg for 10 min, brake on.
11) Pour supernatant into a new standard tube by holding the SepMate™ tube in a vertical position.
12) Top up the tube to 50 mL with recommended medium to wash enriched cells.
13) Centrifuge 300 xg for 10 min, brake off.
14) Discard supernatant by aspiration, there will be some liquid remaining, agitate/flick it to break up the cell pellet.
15) Repeat steps 12 and 13. Remove supernatant by aspiration gently.
16) Resuspend cell pellet in 200 µL of DPBS and 0.2 µL of 1 mg/mL Hoechst dye (1:1000 dilution) using a low binding pipette tip.
17) Transfer contents of tube to a poly-L-lysine-coated well of a 96 well plate. Incubate the plate in the dark (at 37°C and 5% $CO_2$) for 1 hr before visualization.

### Appendix 4. Protocol for Parsortix® PR1

**Preparation of spiked blood sample**

1) Collect approx. 6 mL healthy donor blood into a 10 mL EDTA tube (BD-Plymouth. PL6 7BP.UK, 8083925).
2) Wash out the anticoagulant in another 10 mL EDTA tube by rinsing it with deionised water.
3) Using a low binding pipette tip, add 100 µL of H1975 cell suspension stained with CellTracker™ Green (1000 cells/mL) into this rinsed out 10 mL tube.
4) Pre-coat one serological pipette with 1% BSA and transfer 5 mL blood into this rinsed out tube (containing spiked-in cells).
5) Let it incubate for 30 min in the dark at room temperature and then invert tube 5 times.
6) Add in 1 mL DPBS to the spiked blood sample and invert the spiked blood sample 5 times before processing. This DPBS is added to ensure the blood sample runs smoothly through the PR1.

**Priming the cassette**

1) Prime the cassette with ethanol by choosing program PX2_PF and pressing 'RUN'. Press 'START'.
2) Prompt will say 'Insert new cassette' - press 'OK'.

Machine will beep when finished after a few min- press 'OK' and then 'Continue'.

**Blood Processing**

1) Choose program PX2_S99F and press 'RUN' and then press 'START'.
2) Prompt will say 'rinse vacutainer' – Don't click OK, put in an empty tube and then click 'OK'.
3) Prompt will say 'attach vacutainer'. Put in the spiked blood sample tube and then click 'OK'.
4) The sample should start to fill the cassette within about 30 seconds. If the following prompt appears 'preparing sample', press 'OK'.
5) Flick the sample tube every 20 min.
6) Machine will say finished S99F – Press 'OK' and continue.

**Appendix 4.1. Protocol for Parsortix® PR1 in-cassette staining**

1) Leave the separation cassette in the clamping mechanism. Prepare the following reagents:
    a. **<u>Fixative:</u>** Add 360 µL DPBS and 40 µL formaldehyde solution (Sigma-Aldrich, F8775) to a Corning 15 mL tube.
    b. **<u>Antibody cocktail:</u>** To another Corning 15 mL tube, add the following: 3.5 µL CD45 antibody (1:120 dilution) (35-Z6, Alexa Fluor 647, SantaCruz Biotechnology, sc-1178), 3.5 µL pan-Cytokeratin antibody (1:120 dilution) (C11, Alexa Fluor 488, Biotechnology, sc-8018), 3.5 µL Cytokeratin 19 antibody (1:120 dilution) (A53-B/A2, Alexa Fluor 488, SantaCruz Biotechnology, sc-6278), 3.5 µL EpCAM antibody (1:120 dilution) (HEA125, Alexa Fluor 488, SantaCruz Biotechnology, sc-59906), 3.5 µL Vimentin antibody (1:120 dilution) (V9, Alexa Fluor 546, SantaCruz Biotechnology sc-6260), 3.5 µL 1mg/mL Hoechst 33342 dye and 399 µL of permeabilization buffer Inside Perm (MACS Miltenyi Biotec, Inside stain Kit, order no 130-090-477). This recipe was based on our optimised antibody protocol.
2) Add the correct tubes to the correct lines, ensuring that the end of the tubing reaches the bottom of the tube and is immersed in the fluid.
3) Place a foil tray over the cassette to protect it from the light.
4) Select PX2_stain 3 and press 'RUN'.
5) Add separation cassette and click 'OK'.
6) At prompt 'Fix + Perm line 1', click 'OK'.
7) At prompt 'Abs + DAPI line 2', click 'OK'.

**Appendix 4.2 Protocol for Parsortix® PR1 cell harvest**

1) Do Pre-harvest Flush.

2) At prompt “Insert cleaning cassette”
3) Open the clamp, remove the separation cassette, examine under microscope if intended and keep safe.
4) Insert the cleaning cassette and press [OK].
5) At prompt “Empty rgt tubes”. Ensure reagent tubes are empty. Press [OK] to start the flush process.
6) Select the protocol PX2_CT2 (Cell harvesting protocol) and press [Run] then [Start].
7) At prompt “Finished CT2” press [OK] then [Continue] to return to the main menu on the screen.
8) Remove the cleaning cassette and reinsert the separation cassette.
9) Ensure the harvest waste tube is empty.
10) In the main menu screen, select the protocol PX2_H and press [Run] then [Start]. When prompted, rotate the harvest valve anticlockwise to the position HAR and press [OK].
11) At prompt “Start”. Remove the harvest line from the harvest waste tube and clean it with an alcohol-soaked wipe. Allow it to dry.
12) Place a low binding Eppendorf tube beneath the harvest line.
13) Press [OK] to start the harvest. A volume of approx. 200 µl will flow through the harvest line. At prompt “Further flush?” Press [No].
14) Place the harvest line back in the harvest waste tube.
15) On prompt, rotate the harvest valve clockwise to the position “SEP” and press [OK].
16) At prompt “Finished H” press [OK] then [Continue] to return to the main menu on the screen
17) Add 0.2 µL of 1mg/mL of Hoechst dye to volume containing harvested cells in the Eppendorf tube.
18) Pipette up and down using low binding pipette tip.
19) Transfer contents of Eppendorf tube to a poly-L-lysine-coated well of a 96 well plate. Incubate the plate in the dark (at 37°C and 5% $CO_2$) for 1 hr before visualization.

### Appendix 5. Protocol for Parsortix® Plus (PX+)

#### Appendix 5.1. Protocol for Parsortix® Plus (PX+) cell harvest

1) Choose PXP_SMP_v1 protocol. Press [Run] and when prompted press [Start].
2) Open the sample door and when prompted “Place blood in sample holder”, insert the spiked blood sample into the tube cube and click [OK].
3) When prompted, close the sample door and open the viewing window.
4) The sample mount will begin to rise and when prompted “Dip tubes to make contact with sample?”, ensure the dip tubes will make contact inside the vacutainer by observing through the viewing window. If this is the case, click [YES].
5) When prompted “Dip tubes central / complete mounting?”, click [YES] and the sample will complete mounting.
6) Close the viewing window when prompted and click [OK].

7) When prompted “Insert fresh waste bottle”, open the harvest drawer and insert an empty 4 mL Nalgene bottle in the waste compartment. Click [OK].
8) When prompted “Insert fresh harvest vessel/well”, add a new 1.5 mL low binding Eppendorf into the harvest compartment and click [OK]
9) Close the harvest door and when prompted “Prepare Harvest / Keep Door Open?”, click [NO] to continue with the protocol.
10) When prompted “Preparing / Open Cassette…”, click [OPEN] to open the cassette clamp.
11) The user can dispose of any cassette already in the clamp and insert a new separation cassette. Close the cassette clamp.
12) Once the separation has finished open the sample door and remove the sample tube and click [OK].
13) When prompted “Insert clean wash tube”, insert an empty vacutainer to the tube cube and click [OK].
14) When prompted, close the sample door, and open the viewing window.
15) The sample mount will begin to rise and when prompted “Dip tubes to make contact with sample?”, ensure the dip tubes will make contact inside the vacutainer by observing through the viewing window. If this is the case, click [YES] and the sample mount will complete mounting. -
16) When prompted “Dip tubes central / complete mounting?”, click [YES] and the sample will complete mounting.
17) When prompted, close the viewing window, and click [OK].
18) The user will then be prompted “Count capture in cassette?”. - click [NO] to continue to harvest.
19) Once the captured CTCs have been harvested, the harvest door will open, and the user prompted “Finishing Harvesting / Take Harvest Vessel”. Remove the 1.5 mL Eppendorf that contains the harvested cells.
20) Once the harvest has been taken, close the harvest door, and continue to the clean section by clicking [YES] when prompted.
21) Add 0.2 µL of 1mg/mL of Hoechst 33342 dye to the volume containing harvested cells in the respective Eppendorf tube.
22) Pipette up and down using low binding pipette tip.
23) Transfer contents of Eppendorf tube to a poly-l-lysine-coated well of a 96 well plate. Incubate the plate in the dark (at 37°C and 5% $CO_2$) for 1 hr before visualization.

**Appendix 5.2. Protocol for Parsortix® Plus (PX+) in-cassette staining**

As the PX+ does not enable in-cassette staining, at step 18 (Appendix 5.1), click [Yes] to the following prompt : ‘’Count capture in cassette?’’. Remove the cassette and place it on the PR1 for in-casette staining (Appendix 4.1.).
