## Supplemental Fig 1-2 for "A comparative study of circulating tumor cell isolation and enumeration technologies in lung cancer"

### Supplementary Figures

**Fig. S1.** Automated imaging protocol for scanning a well of 96-well plate at 4X to image cell harvest from CellMag™, EasySep™, RosetteSep™, Parsortix® PR1 and PX+ systems and the control wells.

**Fig. S2.** Automated imaging protocol for scanning the whole Parsortix® slides at 4X to image after Parsortix® PR1 and PX+ in-cassette staining process.

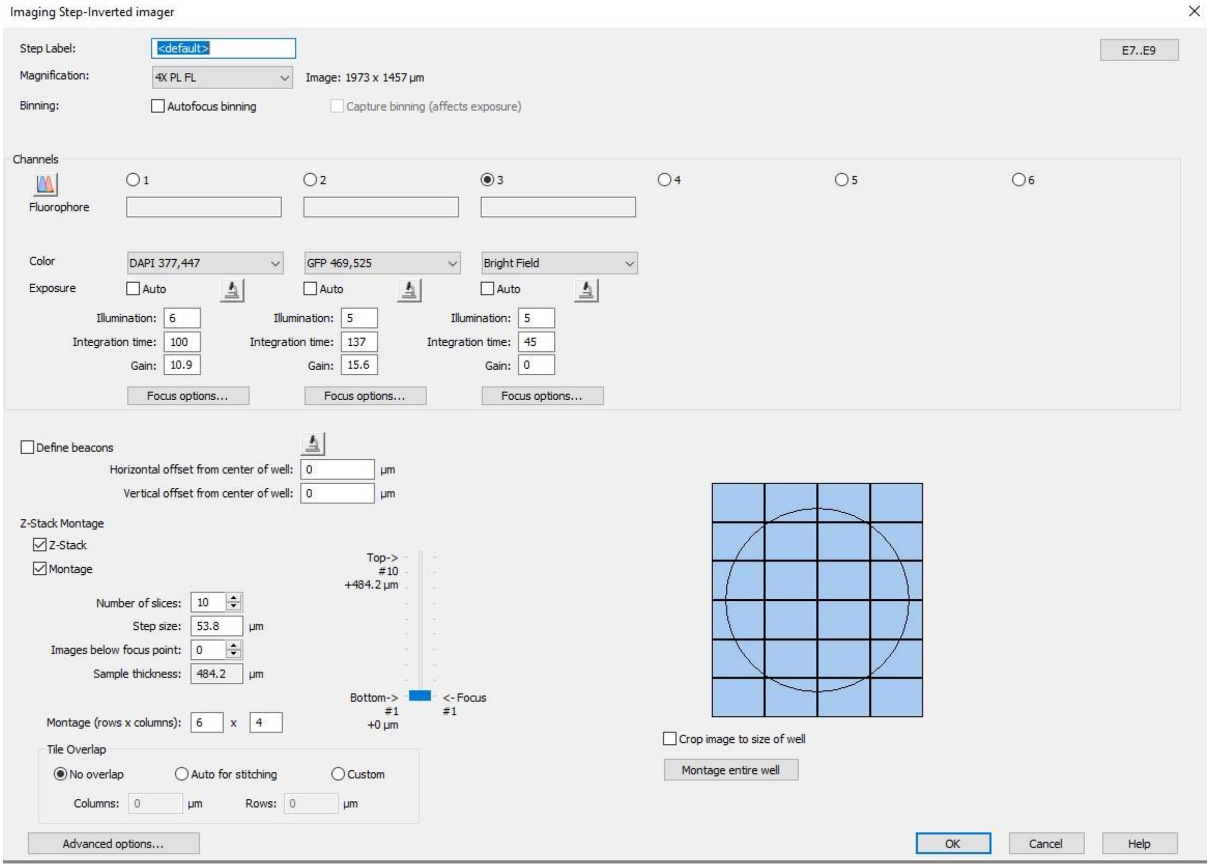


***Fig. S1. Automated imaging protocol for scanning a well of 96-well plate at 4X to image cell harvest from CellMag™, EasySep™, RosetteSep™, Parsortix® PR1 and PX+ systems and the control wells.*** *The recovered cells (in the 96 well plate) from the CellMag™, EasySep™, RosetteSep™, PR1 and PX+ (Harvest) systems and their respective control wells were imaged using an automated imaging protocol and the BioTek Lionheart FX automated microscope. The above protocol (Gen5 3.12 software) was established which can take images (tiles) from all areas of the well at a magnification of 4X. In this protocol, we divided the well into 6x4 image 'tiles' which were then 'stitched together' to create a 'montage' of the entire well. These images were taken using a Z-stack (10 focal planes). This protocol was set up to take images with the DAPI (Hoechst), GFP (CellTracker Green) and Bright Field channels. The Lionheart FX protocol file can be found on FigShare under the DOI:* ***http://doi.org/10.6084/m9.figshare.25146299***

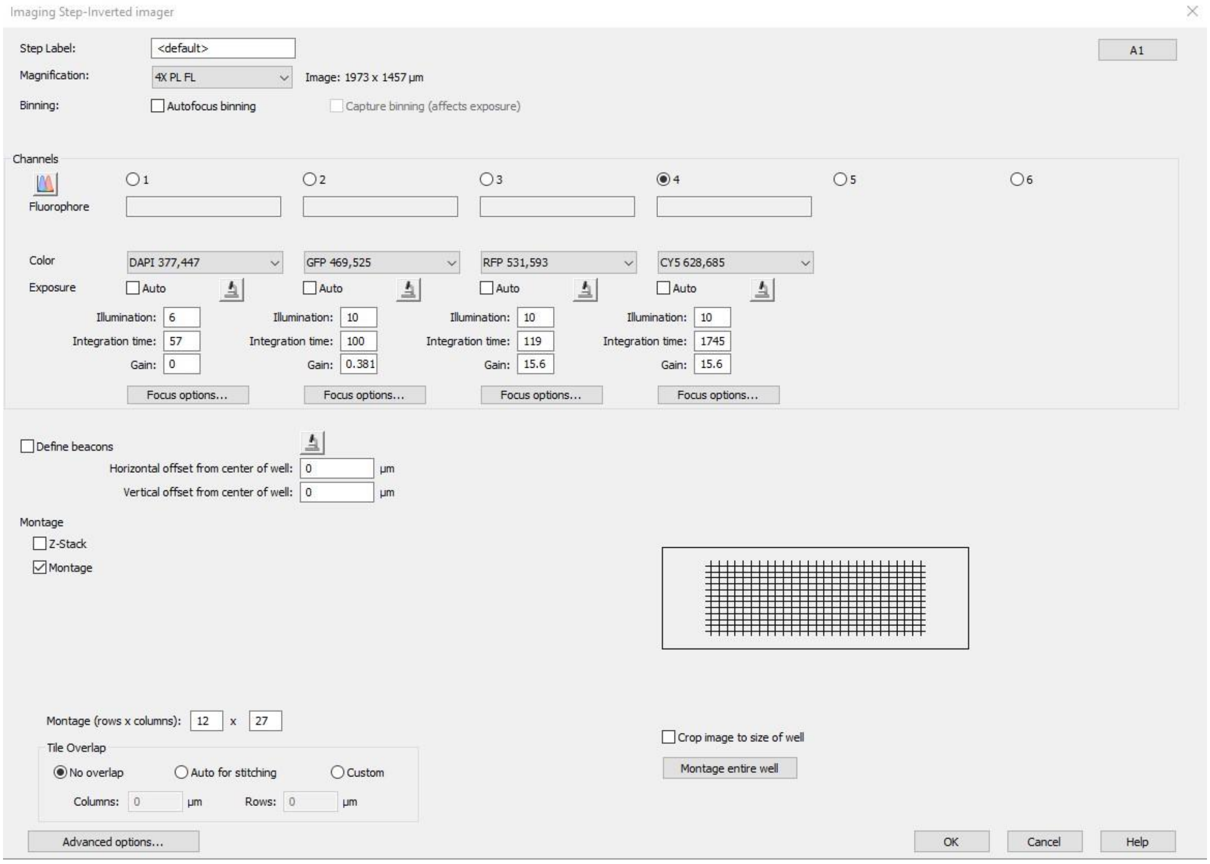


***Fig. S2. Automated imaging protocol for scanning the whole Parsortix® slides at 4X to image after Parsortix® PR1 and PX+ in-cassette staining process.*** *From the PR1 and PX+ in-cassette staining experiments, the spiked cells captured and stained on the Parsortix® slides were visualised using an automated imaging protocol on the BioTek Lionheart FX automated microscope and Gen5 3.12 software. The above protocol was used to scan the whole slide at 4X with the DAPI (Hoechst), GFP (CellTracker™ Green, CK, EpCAM), RFP (Vimentin), CY5 (CD45) channels. The slide was divided into 12x27 (rows x columns) image 'tiles' which were then 'stitched together' to create a 'montage' of the entire slide. The Lionheart FX protocol file can be found on FigShare under the DOI:* ***http://doi.org/10.6084/m9.figshare.25146248***
