## Supplemental Table 1-3 for "A comparative study of circulating tumor cell isolation and enumeration technologies in lung cancer"

### Supplementary Tables

**Table S1.** Overview of the percentage of recovery rates from CTC isolation technologies tested in this study.

**Table S2.** Results of Tukey's multiple comparisons test displaying the difference and significance in recovery rates (**normalized to control**).

**Table S3.** Results of Tukey's multiple comparisons test displaying the difference and significance in recovery rates (**not normalized to control**).

**Table S1.** Overview of the percentage of recovery rates from CTC isolation technologies tested in this study.

| Replicate | CellMag™ | EasySep™ | RosetteSep™ | PR1 Harvest | PX Harvest | PR1 In-cassette | PX+ In-cassette |
|---|---|---|---|---|---|---|---|
| 1 | 53.59 | 9.47 | 16.57 | 37.89 | 33.68 | 47.62 | 43.81 |
| 2 | 76.18 | 19.2 | 22.69 | 32.67 | 26.73 | 49.52 | 54.29 |
| 3 | 80.48 | 24.75 | 36 | 48.65 | 45.95 | 50.94 | 25.47 |
| **Mean** | **70.08** | **17.81** | **25.09** | **39.74** | **35.45** | **49.36** | **41.19** |
| **SD** | **14.44** | **7.73** | **9.93** | **8.15** | **9.73** | **1.67** | **14.59** |

**Table S2.** Results of Tukey's multiple comparisons test displaying the difference and significance in recovery rates (**normalized to control**)[a]

| Tukey's multiple comparisons test | Mean Diff. | 95.00% CI of diff. | Significant? | Summary | Adjusted P Value |
|---|---|---|---|---|---|
| CellMag™ vs. EasySep™ | 52.28 | 23.54 to 81.02 | Yes | *** | 0.0004 |
| CellMag ™vs. RosetteSep™ | 45.00 | 16.26 to 73.74 | Yes | ** | 0.0015 |
| CellMag™ vs. PR1 Harvest | 30.35 | 1.607 to 59.09 | Yes | * | 0.0355 |
| CellMag™ vs. PX Harvest | 34.63 | 5.890 to 63.37 | Yes | * | 0.0141 |
| CellMag ™ vs. PR1 In-cassette | 20.72 | -8.016 to 49.46 | No | ns | 0.2439 |
| CellMag™ vs. PX+ In-cassette | 28.89 | 0.1536 to 57.63 | Yes | * | 0.0484 |
| EasySep ™ vs. RosetteSep™ | -7.280 | -36.02 to 21.46 | No | ns | 0.9723 |
| EasySep™ vs. PR1 Harvest | -21.93 | -50.67 to 6.810 | No | ns | 0.1961 |
| EasySep™ vs. PX Harvest | -17.65 | -46.39 to 11.09 | No | ns | 0.4043 |
| EasySep™ vs. PR1 In-cassette | -31.55 | -60.29 to -2.814 | Yes | * | 0.0274 |
| EasySep™ vs. PX+ In-cassette | -23.38 | -52.12 to 5.356 | No | ns | 0.1489 |
| RosetteSep™ vs. PR1 Harvest | -14.65 | -43.39 to 14.09 | No | ns | 0.6030 |
| RosetteSep™ vs. PX Harvest | -10.37 | -39.11 to 18.37 | No | ns | 0.8705 |
| RosetteSep™ vs. PR1 In-cassette | -24.27 | -53.01 to 4.466 | No | ns | 0.1251 |
| RosetteSep™ vs. PX+ In-cassette | -16.10 | -44.84 to 12.64 | No | ns | 0.5033 |
| PR1 Harvest vs. PX Harvest | 4.283 | -24.46 to 33.02 | No | ns | 0.9983 |
| PR1 Harvest vs. PR1 In-cassette | -9.623 | -38.36 to 19.12 | No | ns | 0.9037 |
| PR1 Harvest vs. PX+ In-cassette | -1.453 | -30.19 to 27.29 | No | ns | >0.9999 |
| PX Harvest vs. PR1 In-cassette | -13.91 | -42.65 to 14.83 | No | ns | 0.6546 |
| PX Harvest vs. PX+ In-cassette | -5.737 | -34.48 to 23.00 | No | ns | 0.9916 |
| PR1 In-cassette vs. PX+ In-cassette | 8.170 | -20.57 to 36.91 | No | ns | 0.9526 |

*a Analysis was performed using GraphPad Prism (version 8.0.2).*

**Table S3.** Results of Tukey's multiple comparisons test displaying the difference and significance in recovery rates (**not normalized to control**)[a]

| Tukey's multiple comparisons test | Mean Diff. | 95.00% CI of diff. | Significant? | Summary | Adjusted P Value |
|---|---|---|---|---|---|
| CellMag™ vs. EasySep™ | 77.33 | 31.81 to 122.9 | Yes | *** | 0.0007 |
| CellMag™ vs. RosetteSep™ | 67.00 | 21.48 to 112.5 | Yes | ** | 0.0027 |
| CellMag™ vs. PR1 Harvest | 49.33 | 3.810 to 94.86 | Yes | * | 0.0299 |
| CellMag™ vs. PX Harvest | 54.00 | 8.476 to 99.52 | Yes | * | 0.0158 |
| CellMag™ vs. PR1 In-cassette | 44.33 | -1.190 to 89.86 | No | ns | 0.0586 |
| CellMag™ vs. PX+ In-cassette | 53.00 | 7.476 to 98.52 | Yes | * | 0.0182 |
| EasySep™ vs. RosetteSep™ | -10.33 | -55.86 to 35.19 | No | ns | 0.9839 |
| EasySep™ vs. PR1 Harvest | -28.00 | -73.52 to 17.52 | No | ns | 0.4025 |
| EasySep™ vs. PX Harvest | -23.33 | -68.86 to 22.19 | No | ns | 0.5974 |
| EasySep™ vs. PR1 In-cassette | -33.00 | -78.52 to 12.52 | No | ns | 0.2392 |
| EasySep™ vs. PX+ In-cassette | -24.33 | -69.86 to 21.19 | No | ns | 0.5537 |
| RosetteSep™ vs. PR1 Harvest | -17.67 | -63.19 to 27.86 | No | ns | 0.8297 |
| RosetteSep™ vs. PX Harvest | -13.00 | -58.52 to 32.52 | No | ns | 0.9516 |
| RosetteSep™ vs. PR1 In-cassette | -22.67 | -68.19 to 22.86 | No | ns | 0.6267 |
| RosetteSep™ vs. PX+ In-cassette | -14.00 | -59.52 to 31.52 | No | ns | 0.9327 |
| PR1 Harvest vs. PX Harvest | 4.667 | -40.86 to 50.19 | No | ns | 0.9998 |
| PR1 Harvest vs. PR1 In-cassette | -5.000 | -50.52 to 40.52 | No | ns | 0.9997 |
| PR1 Harvest vs. PX+ In-cassette | 3.667 | -41.86 to 49.19 | No | ns | >0.9999 |
| PX Harvest vs. PR1 In-cassette | -9.667 | -55.19 to 35.86 | No | ns | 0.9885 |
| PX Harvest vs. PX+ In-cassette | -1.000 | -46.52 to 44.52 | No | ns | >0.9999 |
| PR1 In-cassette vs. PX+ In-cassette | 8.667 | -36.86 to 54.19 | No | ns | 0.9935 |

*a Analysis was performed using GraphPad Prism (version 8.0.2).*
